# Mechanistic insights into gene expression changes and electric organ discharge elongation in mormyrid electric fish

**DOI:** 10.1101/2024.09.02.610879

**Authors:** Mauricio Losilla, Jason R. Gallant

## Abstract

Electric organ discharge (EOD) duration in African weakly electric fish (Mormyridae) is the most variable waveform component between species and the basis for distinguishing species-specific signals. EOD duration is thought to be influenced by morphological and physiological features of electrocytes (the cells that comprise the electric organ), but the mechanistic details are poorly understood. It has long been known that EOD duration is modulated by androgen hormones, affording an opportunity to identify gene expression correlates of EOD duration differences. We induced EOD elongation in the mormyrid *Brienomyrus brachyistius* by administering 17α-methyltestosterone (17αMT) to three treatment groups: control (no 17αMT exposure), T1day and T8day (samples taken one and eight days after a single exposure to 17αMT, respectively). We then performed RNAseq, differential gene expression, and functional enrichment analysis to detect gene expression changes during EOD duration change. Our analyses indicate 44 genes whose expression changed in tandem with EOD elongation and include genes responsible for actin filaments and microtubules, extracellular matrix organization, and membrane lipid metabolism. Additionally, we found expression changes in one Na^+^ channel beta subunit, and five K^+^ voltage-gated channels. Together, these genes point toward specific cellular processes that contribute to morphological and physiological changes that contribute to EOD duration changes.

## Introduction

Despite little ecological differentiation, African weakly electric fish (Mormyridae) hold one of the highest rates of speciation among ray-finned fishes (Carlson et al. 2011; Rabosky et al. 2013). It is suspected that a crucial factor of their extraordinary diversification is divergence in their electric organ discharges (EODs). In Paramormyrops, EODs have evolved faster than size, morphology, and trophic ecology (Arnegard et al. 2010a). These electric signals, crucial to electrolocation (Lissmann and Machin 1958; von der Emde et al. 2008) and communication (Möhres 1957; Kramer 1974), are effective mechanisms of prezygotic isolation (Hopkins and Bass 1981; Arnegard et al. 2006), and often are the most reliable method for identifying species (Hopkins 1981; Bass 1986; Arnegard et al. 2005).

Species-specific mormyrid EOD waveforms vary primarily in their complexity (number of phases), polarity (alterations to waveform shape caused by large a P0), and duration (Hopkins 1999b; Sullivan et al. 2002), the latter being the most variable factor between species (Arnegard and Hopkins 2003; Gallant et al. 2011) and the basis for recognition of species-specific signals (Hopkins and Bass 1981; Arnegard et al. 2006). EOD duration differs up to 100x among species (Hopkins 1999a), ranging from 0.2 to > 15 ms (Lavoué et al. 2008); it is often sexually dimorphic (Bass and Hopkins 1983; Hopkins 1999a), with differences between male and female adult EODs often exacerbated during the rainy season breeding period (Hopkins and Bass 1981).

The distinctive features of the EOD waveform are a consequence of the physiological, morphological, and ultrastructural characteristics of the electrocytes (Carlson 2002). Mormyrid EODs are produced by the synchronized discharge of the electrocytes, the constitutive cells of the electric organ (EO), located in the caudal peduncle (Hopkins 1986). The EOD from an individual electrocyte is composed of the sum of the action potentials (APs) from different parts of its excitable membrane. The two largest phases of a typical mormyrid EOD, called P1 and P2, are generated by the combined APs of the posterior and anterior faces of the electrocyte membrane (Bennett and Grundfest 1961). These two membranes are thought to receive the depolarization signal simultaneously, but ions begin flowing through the posterior face first (Stoddard and Markham 2008). The reasons for this delay could be physiological (ion channels) and/or due to greater capacitance of the more intricately folded anterior face; however, the majority of the two APs still overlap. Therefore, the resulting EOD is the net current of overlapping in-and outgoing ionic currents at the two faces of the membrane (reviewed in (Markham 2013; Dunlap et al. 2017), see Stoddard and Markham (2008) for a simplified figure of this model). This model is thought to apply broadly to biphasic EODs from mormyrids and the Gymnotiformes (a diverse clade of independently evolved neotropical weakly electric fish). Biphasic EODs have only P1 and P2, but several mormyrids produce triphasic waveforms with an additional, initial phase, P0. Their electrocytes have stalks that penetrate the electrocyte (Pa configuration), P0 is produced when current flows through these penetrating stalks (more details in (Bennett and Grundfest 1961; Szabo 1961; Bennett 1971)).

EOD duration is thought to be regulated by both morphological and physiological aspects. Electrocytes with a larger membrane surface area have been correlated with EODs of longer duration (Bennett 1971; Paul et al. 2015), especially when the area increase is more prominent in the anterior face of the membrane (Bass et al. 1986; Freedman et al. 1989; Paul et al. 2015; Nguyen et al. 2020). Presumably, the larger area raises the capacitance of the membrane, and thus increases the time it takes to start depolarizing (Bass et al. 1986; Bass and Volman 1987; Stoddard and Markham 2008). If this delay is more pronounced in the anterior face of the membrane, the net effect should be to extend EOD duration. In addition, changes to the abundance or kinetics of the ion channels in the electrocyte membrane are expected to also modulate EOD duration. In gymnotiforms, it has been demonstrated that the main electrocyte ionic currents are generated by voltage-gated Na^+^ and K^+^ currents (e.g. (Shenkel and Sigworth 1991; Ferrari and Zakon 1993)) and it is broadly expected that the same holds true in mormyrid electrocytes, where both Na^+^ (Zakon et al. 2006; Arnegard et al. 2010b) and K^+^ (Nagel et al. 2017; Swapna et al. 2018) voltage-gated channels are expressed, and differential expression of ion channel genes has been linked with interspecific differences in EOD waveform (Lamanna et al. 2015; Losilla et al. 2020).

We recently performed an examination of differential gene expression among closely related members of the genus *Paramormyrops* (Losilla et al. 2020) where we identified expression correlates of EOD duration. Based on the results of this study, we were motivated to link observed patterns of differential gene expression between species to mechanisms underlying changes in EOD duration that occur within the lifetime of an individual. Previous studies have indicated that androgen hormones, including 17α-methyltestosterone (17αMT) increase EOD duration in several mormyrid species, an effect that mimics natural sex differences (Bass and Hopkins 1983, 1985; Bass et al. 1986; Bass and Volman 1987; Freedman et al. 1989; Herfeld and Moller 1998). Although elongated EODs during the breeding season are a male character, androgen hormone treatment of female mormyrids elicits a male-like response in the duration of the EOD (Bass and Hopkins 1983, 1985; Bass et al. 1986; Bass and Volman 1987; Freedman et al. 1989; Herfeld and Moller 1998). Androgen hormones provoke similar responses on the duration of male and female EODs in gymnotiforms (e.g. (Mills and Zakon 1987)).

Treatment with androgen hormones can provoke large changes in EOD duration under controlled circumstances, providing a hitherto unexplored opportunity to identify the expression correlates of EOD duration differences. In this study, we leveraged this paradigm to investigate the molecular underpinnings of changes in EOD duration in the mormyrid *Brienomyrus brachyistius*, a species where males greatly elongate their EOD under breeding conditions, influenced by their social status and in connection with androgen levels in blood (Carlson et al. 2000). We experimentally applied 17αMT to *B. brachyistius* individuals over a 9-day experimental period and examined patterns of differential gene expression over the course of this treatment compared to control organisms. Unlike our previous study, which compared gene expression between species, this analysis focused on conspecific adult individuals in controlled environments, eliminating critical confounding factors like phylogenetic divergence and additional EOD variation. Our analysis strongly supports known aspects of morphological and physiological bases of EOD duration, and for the first time identifies specific genes and broad cellular processes that alter morphological and physiological properties of electrocytes during seasonally plastic EOD changes.

## Methods

### Study fish

Our study was performed on 20 adult (>=110mm) *Brienomyrus brachyistius* purchased through the pet trade. Males were identified by the presence of an indentation in the anal fin, a common dimorphic feature in mormyrids. Sex was confirmed by gonad inspection after which we believe that 19 fish were males and one was female.

### Experimental conditions

Subjects were housed individually in 25 cm width x 50 cm depth x 30 cm tanks filled with 30 L of water and equipped with one mechanical and biological filter. We kept the fishes under 12 h light cycles, fed them blackworms daily, monitored levels of ammonia, nitrites, and nitrates, and performed 30% water changes as needed during acclimation periods (no water changes were carried out after experimental treatments were applied). The following per tank water conditions were monitored daily or every other day and remained within the indicated limits: conductivity 500-570 µS/cm, temperature 24-26 °C, pH 7.0-7.5. Each tank was provided with a custom-built housing arrangement that served as a fish shelter but also facilitated measuring EODs with the fish at a stable location. These arrangements consisted of a PVC tube attached to an aquarium plastic egg crate via two zip ties, the egg crate was cut to the width of the tank and fixed to the bottom with four suction cups with hose clips, each with a short piece of PVC tube acting as a lock. The housing tube laid centered at the bottom, oriented from the front to the back end of the tank (Fig. 1).

**Fig. 1.**
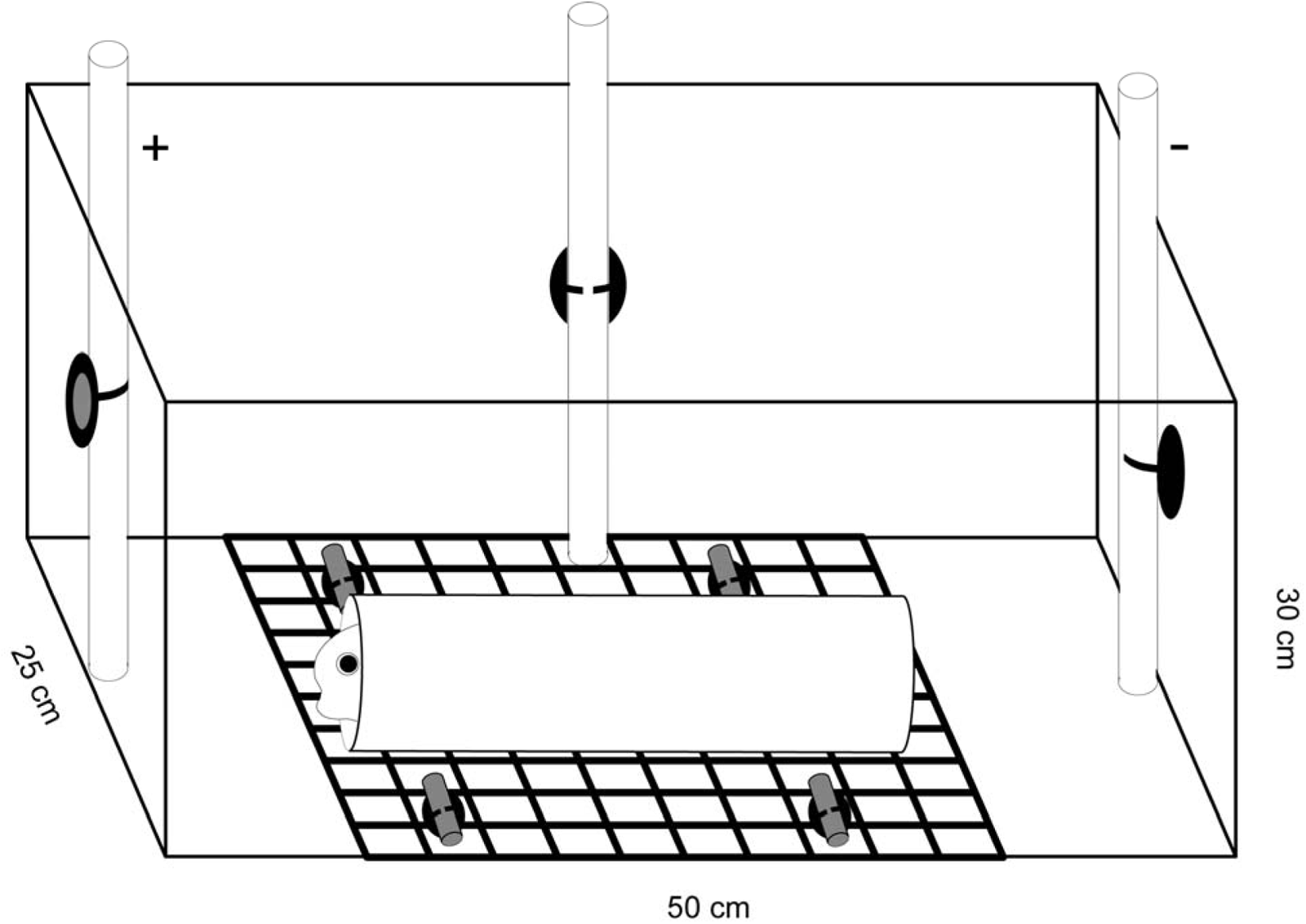
Experimental fish tanks’ configuration, showing the housing arrangement attached to the middle of the tank’s bottom and the fixed electrode locations.

### EOD recordings

For each subject, we recorded daily EOD waveforms (median 19, range 14-20) during acclimation and experimental periods. During recording sessions, electrodes were attached to suction cups with hose clips affixed to the center of the front (positive electrode), back (negative), and left (ground) sides of each tank (Fig. 1). The distance between the positive and negative electrodes was 48 cm. Signals were recorded with bipolar chloride-coated silver wire electrodes, amplified (bandwidth =0.0001-50 kHz, gain = 20) with a bioamplifier (model BMA-200; CWE, Inc), digitized using a multifunction DAQ device (model USB-6216 BNC; NI) and the recording software EODrecorder2 (Terminal Configuration = Differential, sampling rate = 250 KHz, voltage range = +/-5 V, npts = 2048). On each EOD, we used custom code to correct baseline noise, normalize by its peak-to-peak amplitude, center time = 0 at P1, and determine the start and stop points (and hence the duration).

### Experimental treatments

Each subject was acclimated to the experimental setup for a minimum of two weeks before the start of a treatment. We deemed a fish acclimated when the CV of EOD duration was < 5% for five consecutive days. Experimental treatments were conducted in two back-to-back cycles, with 8 and 12 fish, respectively. For each cycle, treatment was applied on day 0, immediately after taking the EOD recordings for that day. Fish were assigned to one of three experimental treatments: two groups received 3 mg/L of 17αMT dissolved in 5ml of ethanol 100% administered directly to the tank water and were allowed to survive 1 day (T1day; n=7) or 8 days (T8day; n=7) post treatment. The third group (Control; n=6) received 5ml of 100% ethanol administered directly to the tank water. We monitored the fish and recorded EODs daily, until and including the assigned dissection date. Our experimental setup (fixed housing arrangement, EOD recordings taken in each fish’s tank, and non-invasive 17αMT addition) was designed to minimize fish manipulation and disturbance. The study individuals did not undergo surgery or injections, and once placed in its experimental tank, a fish was not removed or otherwise handled until its dissection date. All procedures complied with federal and state regulations and were approved by Michigan State University’s Office of Environmental Health & Safety, and Institutional Animal Care & Use Committee.

### EOD Duration Analysis

We tested for differences in EOD duration with a one-way ANOVA at 1) day 0, and 2) the last day of treatment (i.e. day 1 for T1day, day 8 for control and T8day). Where significant results were detected, we examined differences in all pairwise treatment comparisons with the post-hoc Tukey HSD test.

### Tissue dissections, RNA extraction and sequencing

After the final EOD recording session, subjects were euthanized with an overdose of MS-222 (0.7 g MS-222 in 2 L of fish tank water; and 1.4 g of sodium bicarbonate). We skinned and dissected caudal peduncles and stored them in RNA-later (Ambion, Inc) following manufacturer’s instructions until processing. All RNA extraction, library preparation, and sequencing were performed by Genewiz, LLC (South Plainfield, NJ). Total RNA was extracted using Qiagen RNeasy Plus Universal mini kit following manufacturer’s instructions (Qiagen, Hilden, Germany). Extracted RNA samples were quantified using Qubit 2.0 Fluorometer (Life Technologies, Carlsbad, CA, USA) and RNA integrity was checked using Agilent TapeStation 4200 (Agilent Technologies, Palo Alto, CA, USA). RNA poly-A selected sequencing libraries were prepared using the NEBNext Ultra II RNA Library Prep Kit for Illumina using manufacturer’s instructions (NEB, Ipswich, MA, USA). Briefly, mRNAs were first enriched with Oligo(dT) beads. Enriched mRNAs were fragmented for 15 minutes at 94 °C. First strand and second strand cDNAs were subsequently synthesized. cDNA fragments were end repaired and adenylated at 3’ends, and universal adapters were ligated to cDNA fragments, followed by index addition and library enrichment by limited-cycle PCR. The sequencing libraries were validated on the Agilent TapeStation and quantified using Qubit 2.0 Fluorometer as well as by quantitative PCR (KAPA Biosystems, Wilmington, MA, USA). The sequencing libraries were clustered on flow cells. After clustering, the flow cells were loaded on to the Illumina HiSeq instrument (4000 or equivalent) according to manufacturer’s instructions. The samples were sequenced using a 2x150bp Paired End (PE) configuration. Image analysis and base calling were conducted by the HiSeq Control Software (HCS). Raw sequence data (.bcl files) generated from Illumina HiSeq was converted into fastq files and de-multiplexed using Illumina’s bcl2fastq 2.17 software. One mismatch was allowed for index sequence identification.

### Read processing and data exploration

We inspected raw and processed reads with FastQC v0.11.7 (Babraham Bioinformatics) and used Trimmomatic v.0.39 (Bolger et al. 2014) to remove library adaptors, low quality reads, and filter small reads. The succeeding steps were executed using scripts included with Trinity v2.11.0 (Grabherr et al. 2011; Haas et al. 2013). We aligned reads from each specimen to the predicted transcripts of the NCBI-annotated (release 100) *B. brachyistius* genome, with bowtie2 v2.3.4.1 (Langmead and Salzberg 2012). Expression quantification was estimated at the gene level using RSEM v1.3.0 (Li and Dewey 2011), followed by exploration of the data with a gene expression correlation matrix based on Euclidean distances and Pearson’s correlation coefficient (for genes with read counts > 10, Trinity’s default parameters).

### Differential Gene Expression

We found no outliers for gene expression or EOD duration, hence we analyzed samples lumped together in regard to sex and experimental cycle. We identified differentially expressed genes (DEG) between experimental treatments in the three possible pairwise comparisons (control vs T8day, control vs T1day, T1day vs T8day) with edgeR v3.20.9 (Robinson and Oshlack 2010) through a script provided with Trinity. For each comparison (contrast), we conservatively identified differentially expressed genes (significant DEG) as those genes with a minimum expression fold change (FC) of 4 and p-value < 0.001 after FDR correction.

### Functional enrichment analysis

We employed mitch v1.8.0 (Kaspi and Ziemann 2020) to detect sets of genes that exhibit joint up- or downregulation across our three contrasts. As inputs, mitch needs a gene set library and profiled expression data. The latter was the whole set (i.e. not filtered by FC or p-value thresholds) of edgeR differential expression results from each contrast. For the gene set library, we used a *Danio rerio* gmt file with gene ontology (GO) (Ashburner et al. 2000; Carbon et al. 2021) terms from the GO domain Biological Process as gene sets. To generate this file, we employed the script update_GO.sh from GeneSCF-v1.1-p3 (Subhash and Kanduri 2016). Since the gene identifiers in the gmt file are different than those of the edgeR results, mitch requires a third input file that relates these gene identifiers. To create this file, we first identified homologous proteins predicted from the *B. brachyistius* reference genome and those predicted from *Danio rerio* (GRCz11) by blastp (BLAST+v2.11.0 (Camacho et al. 2009)). For each protein, the top hit (e-value ≤1e-10) was used for annotation. Then, we used mygene v1.32.0 (Wu et al. 2013; Xin et al. 2016) to match the *D. rerio* proteins to *D. rerio* genes.

We excluded gene sets with fewer than 20 genes from the mitch analysis (minsetsize option in mitch_calc function); and filtered the enrichment result to gene sets with FDR-corrected p-value < 0.01 and the higher dimensional enrichment score (S) > 0.1, to minimize false positives. To further simplify the analysis interpretation, we employed the tool Visualize from AmiGO 2 (Carbon et al. 2009) to build GO graphs with the enriched GO terms, and based on their hierarchy and connectivity, we manually grouped the GO terms into broad functional categories.

### Differentially expressed genes of highest interest for EOD duration

We used two procedures to identify the most interesting genes in relation to EOD duration from the significant DEG detected by edgeR. First, we started with the gene sets in the broad categories of most interest from the previous step. We further filtered these gene sets to those which show an enrichment change that correlates with the observed changes in EOD duration (see code repo for details). We then compared the genes that make up each of these select gene sets with the significant DEG from each contrast, and highlight the genes common two both sources. Our second procedure consisted on manually selecting additional significant DEG from each contrast based on their annotation. We prioritized genes i) from general functional themes suspected to affect EOD phenotype, and ii) whose expression patterns correlate with the observed changes in EOD duration (i.e., we did not prioritize genes found differentially expressed only in the control vs T1day comparison).

Our experimental design facilitates inferences about gene expression over the amount of time subjects were exposed to 17αMT (Fig. 2B). The comparison control vs T8day informs about broad changes in gene expression over the course of the experiment, whereas control vs T1day exposes early changes in gene expression, and T1day vs T8day uncovers late changes in gene expression. For example, a hypothetical gene set may show little ‘broad’ changes (control vs T8day), but important ‘granular’ changes in opposite directions before and after day 1 (control vs T1day, and T1day vs T8day). We note that we never observed conflicts between ‘broad’ and ‘granular’ expression patterns.

**Fig. 2.**
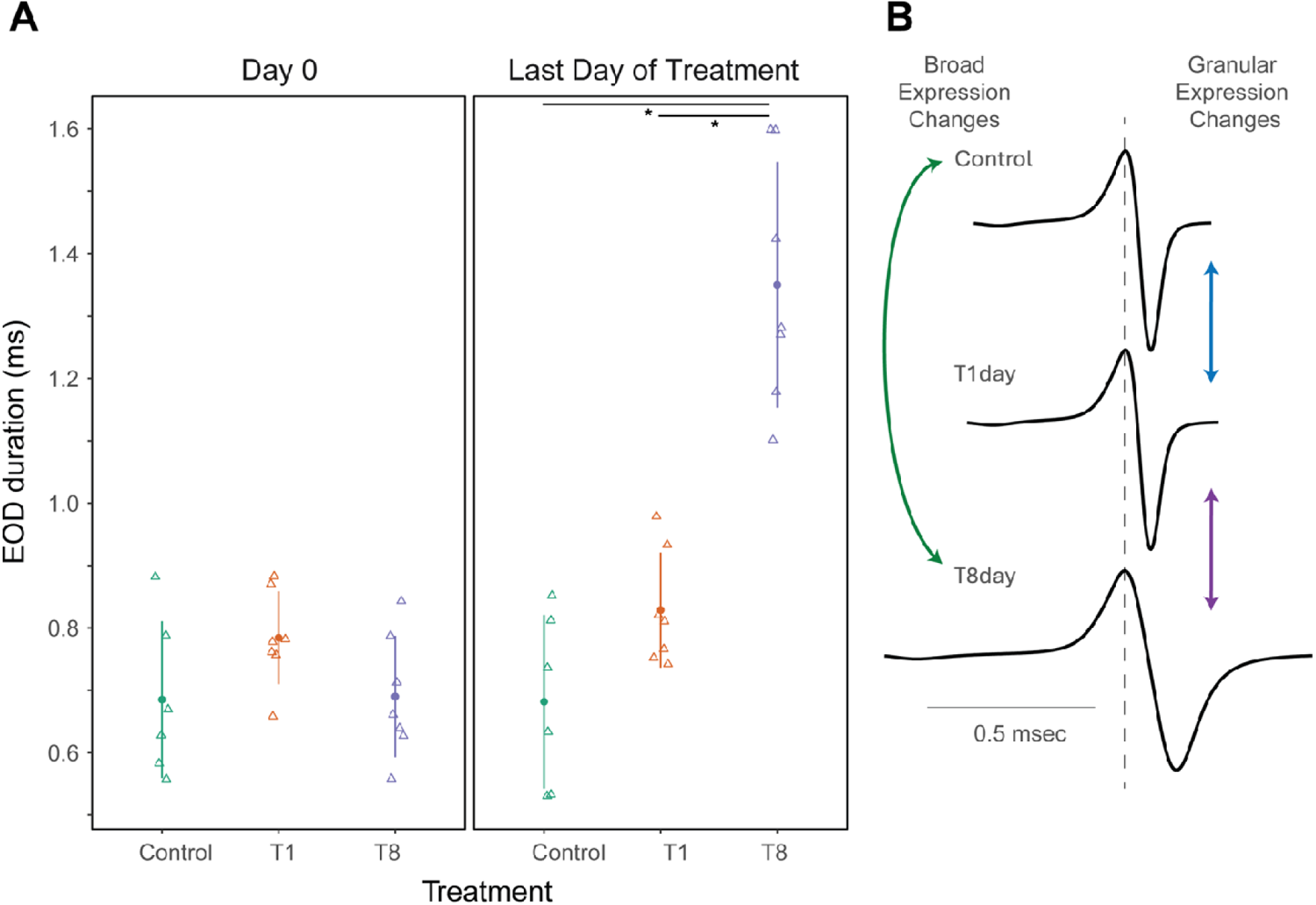
EOD duration in the experimental treatments and classification of gene expression changes. A) EOD duration at the onset (left) and at the end (right) of the experiment. The circle and vertical lines represent the mean EOD duration +/-standard deviation per treatment, open triangles are per fish measurements. A small horizontal jitter was added to better visualize overlapping observations. Each asterisk represents significant differences (p < 0.001) between the treatments connected by its overlying horizontal bar. B) Each graph shows the traces of the averaged EODs from one representative fish per treatment, taken on their last day of recording. The arrows indicate the three pairwise treatment comparisons, and the gene expression changes detected in each comparison are classified as ‘broad’ or ‘granular’, based on each treatment’s time of exposure to 17αMT.

The code used in our analyses is available at https://github.com/msuefishlab/EODduration_geneExpression.

## Results

### EOD duration responses to hormone treatment

There were no differences in the duration of the EOD between the treatments on day 0, immediately before treatment application (p = 0.15, Fig. 2A). On the last day of treatment, shortly before tissue dissection, the duration of the EOD differed between the treatments (p < 0.001); specifically, EOD duration increased in T8day compared to control (p < 0.001) and to T1day (p < 0.001), but it did not differ between control and T1day (p = 0.21) (Fig. 2A). Throughout the experiment, we observed mean EOD duration values stable during the last days of the acclimation period for all treatments and also during the experimental period for treatments control and T1day; however there was a sharp increase in mean EOD duration for fish in the T8day treatment upon addition of 17αMT (Additional file 1). We provide a graphical illustration of the final EOD duration in a representative fish from each treatment in Fig. 2B.

### RNAseq data and differentially expressed genes

We explored the RNAseq data with a heatmap of pairwise correlations of gene expression across all 20 samples. Gene expression patterns across all specimens were highly correlated (Pearson’s r > 0.93), and correlation values were higher between fish from the same treatment (Additional file 2). At the level of treatments, these correlations indicate that the treatments T1day and T8day have a more similar gene expression pattern than with control.

We performed the three possible pairwise differential gene expression (DGE) comparisons between treatments. They each found a similar number of expressed genes (∼20750), and a small percentage (range 0.21 - 1.17%) of DEG (FC > 4, FDR-corrected p-value <0.001). Among the latter, there were always more genes upregulated in the treatment that entailed longer exposures to 17αMT (Table 1). Additional file 3 provides a tabular list of DEGs for every comparison, along with each gene’s annotation, log_2_ FC, p-value, and per fish TMM-normalized expression values.

**Table 1.** All three possible pairwise differential gene expression comparisons with the number of genes expressed, differentially expressed, and their breakdown by treatment.

| comparison | genes expressed | genes differentially expressed (%) | upregulated genes |  |
| --- | --- | --- | --- | --- |
|  |  |  | treatment | total |
| control vs T8day | 20887 | 244 (1.17) | control | 96 |
|  |  |  | T8day | 148 |
| control vs T1day | 20942 | 98 (0.47) | control | 34 |
|  |  |  | T1day | 64 |
| T1day vs T8day | 20401 | 43 (0.21) | T1day | 13 |
|  |  |  | T8day | 30 |

### Functional enrichment analysis

Next, we compared patterns of expression of pre-defined gene sets (GO terms, see methods) across our three pairwise treatment comparisons (contrasts). The level of enrichment (= up- or downregulation) of each gene set in each contrast is quantified by an enrichment score (s, range -1 to 1). These contrasts were constructed such that positive values of s represent upregulation with increased exposure to 17αMT, and negative values of s indicate downregulation with increased exposure to 17αMT.

We detected 96 gene sets enriched across the three contrasts studied (Fig. 3, Additional file 4). To aid with our interpretation, we further grouped these gene sets into 11 broad categories, with 12 gene sets unclassified because they were too general or too isolated in the GO graph (Fig. 3, Additional file 5).

**Fig. 3.**
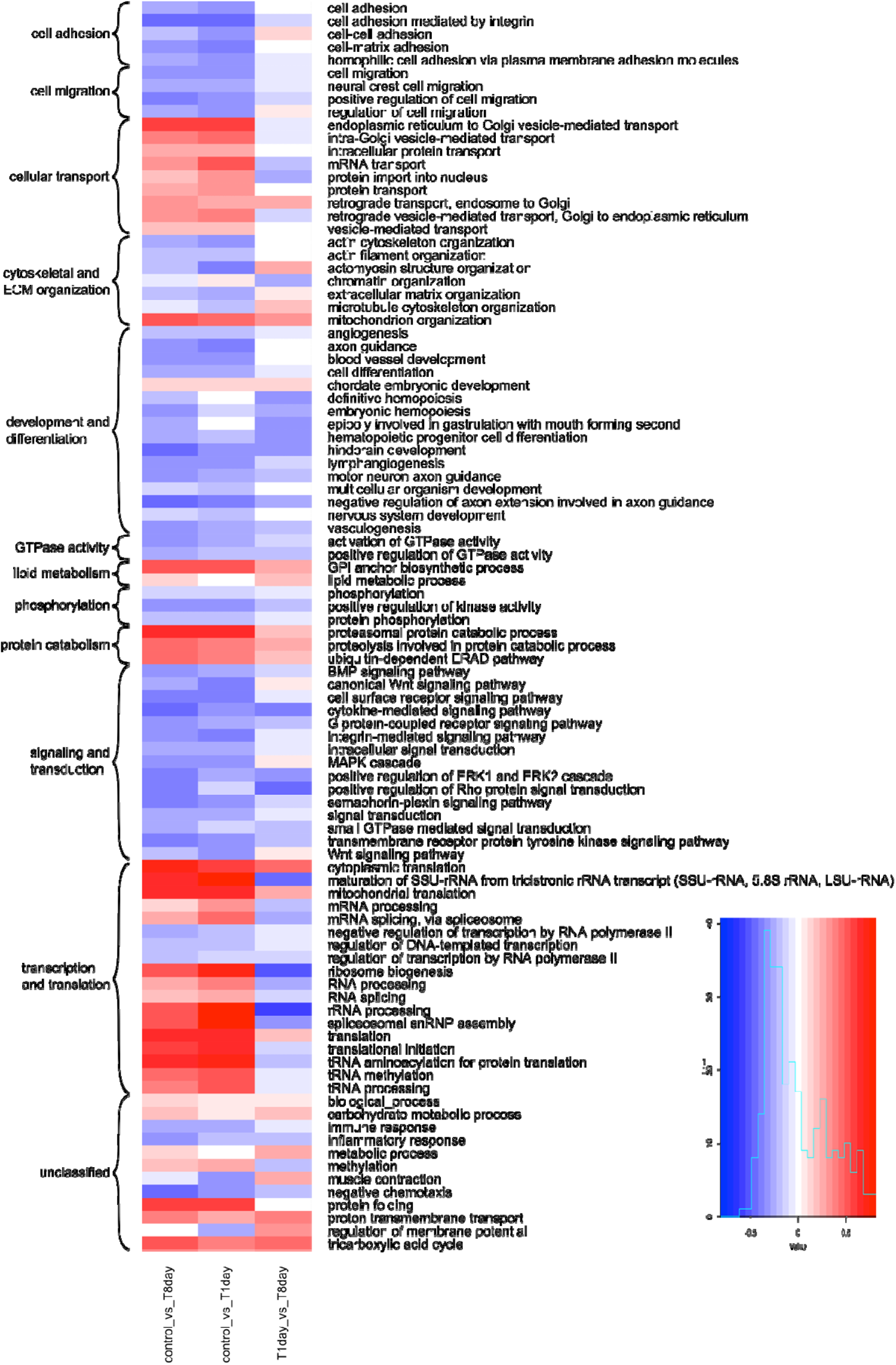
Heatmap of mitch enrichment scores for all significantly enriched gene sets (n = 96, higher dimensional enrichment score > 0.1, FDR p < 0.01) in the three pairwise treatment contrasts studied. Gene sets are further classified into broad categories (curly braces, left).

Overall, the broad enrichment changes (control vs T8day) per gene set are largely driven by early enrichment changes (control vs T1day), whereas later changes in enrichment (T1day vs T8day) are generally more limited. For example, most gene sets in the categories “cell adhesion”, “cell migration”, “phosphorylation”, and “signaling and transduction” were broadly downregulated, and the granular comparisons show that, in most of the constituent gene sets, downregulation happened quickly, while later changes were of smaller magnitude. A similar enrichment pattern, but with mostly upregulated genes instead, is generally observed in the gene sets across the categories “cellular transport”, “protein catabolism”, and “transcription and translation” (Fig. 3).

Because of our overall motivation to examine genes that could contribute to anatomical and physiological changes that would affect EOD duration, we focused on the broad categories of most interest: “cell adhesion”, “cell migration”, “cytoskeletal and ECM organization”, “lipid metabolism”, and the unclassified gene sets “muscle contraction” and “regulation of membrane potential”. We filtered their gene sets to those with an important change in enrichment magnitude and direction before and after day 1 (i.e. between the comparisons control vs T1day and T1day vs T8day), the time when EOD duration started to increase (Additional file 1). We call the resulting 19 gene sets “select gene sets”, they are: “actin cytoskeleton organization”, “actin filament organization”, “actomyosin structure organization”, “cell adhesion”, “cell adhesion mediated by integrin”, “cell migration”, “cell-cell adhesion”, “cell-matrix adhesion”, “chromatin organization”, “extracellular matrix organization”, “GPI anchor biosynthetic process”, “homophilic cell adhesion via plasma membrane adhesion molecules”, “lipid metabolic process”, “microtubule cytoskeleton organization”, “muscle contraction”, “neural crest cell migration”, “positive regulation of cell migration”, “regulation of cell migration”, and “regulation of membrane potential”. We list the genes that comprise the select gene sets in Additional file 4.

### Differentially expressed genes of highest interest for EOD duration

We generated a list of genes that may directly influence EOD duration by i) taking the genes common to at least one select gene set and the DEG from the pairwise comparisons, and ii) from the latter, selecting additional genes that, based on their annotation, could contribute to EOD duration. As a guide, we used the general, functional themes suspected to affect EOD phenotype (Gallant et al. 2012, 2014; Lamanna et al. 2015; Losilla et al. 2020). For simplicity, we sort these genes into the same themes we have used previously (Losilla et al. 2020): “extracellular matrix”, “cation homeostasis”, “lipid metabolism”, and “cytoskeletal & sarcomeric”. We removed from the resulting list those genes found differentially expressed only in the control vs T1day comparison, to prioritize the genes with an expression pattern that resembles the observed changes in EOD phenotype. We also removed the gene *srd5a2* because its product converts testosterone to dihydrotestosterone (DHT). The outcome is the list of DEG of highest interest for EOD duration. We provide the complete list (44 genes), along with functional annotations from UniProt (The UniProt Consortium 2019), in Additional file 6. Select genes are mentioned in the next section and presented in Tables 2-4.

**Table 2.**
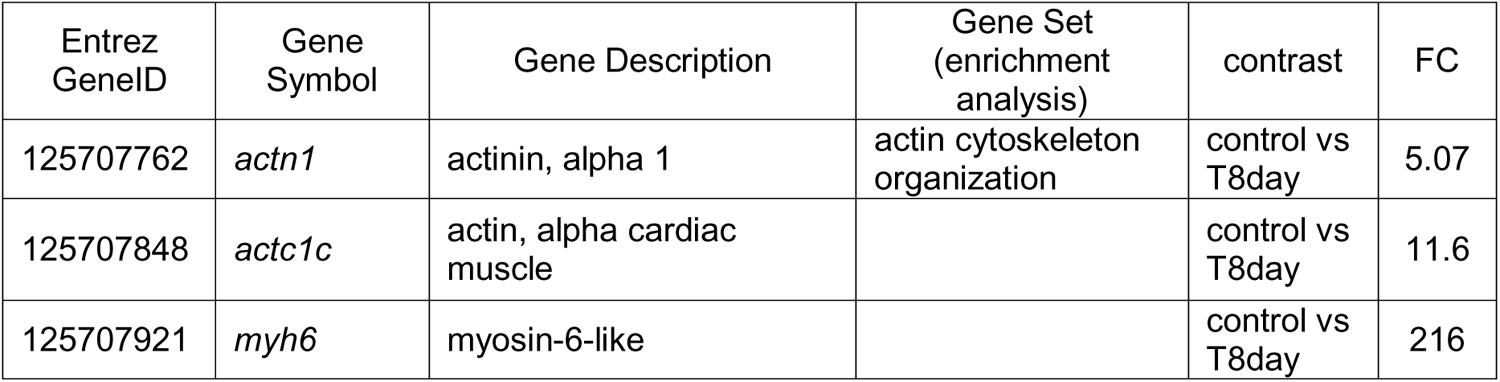
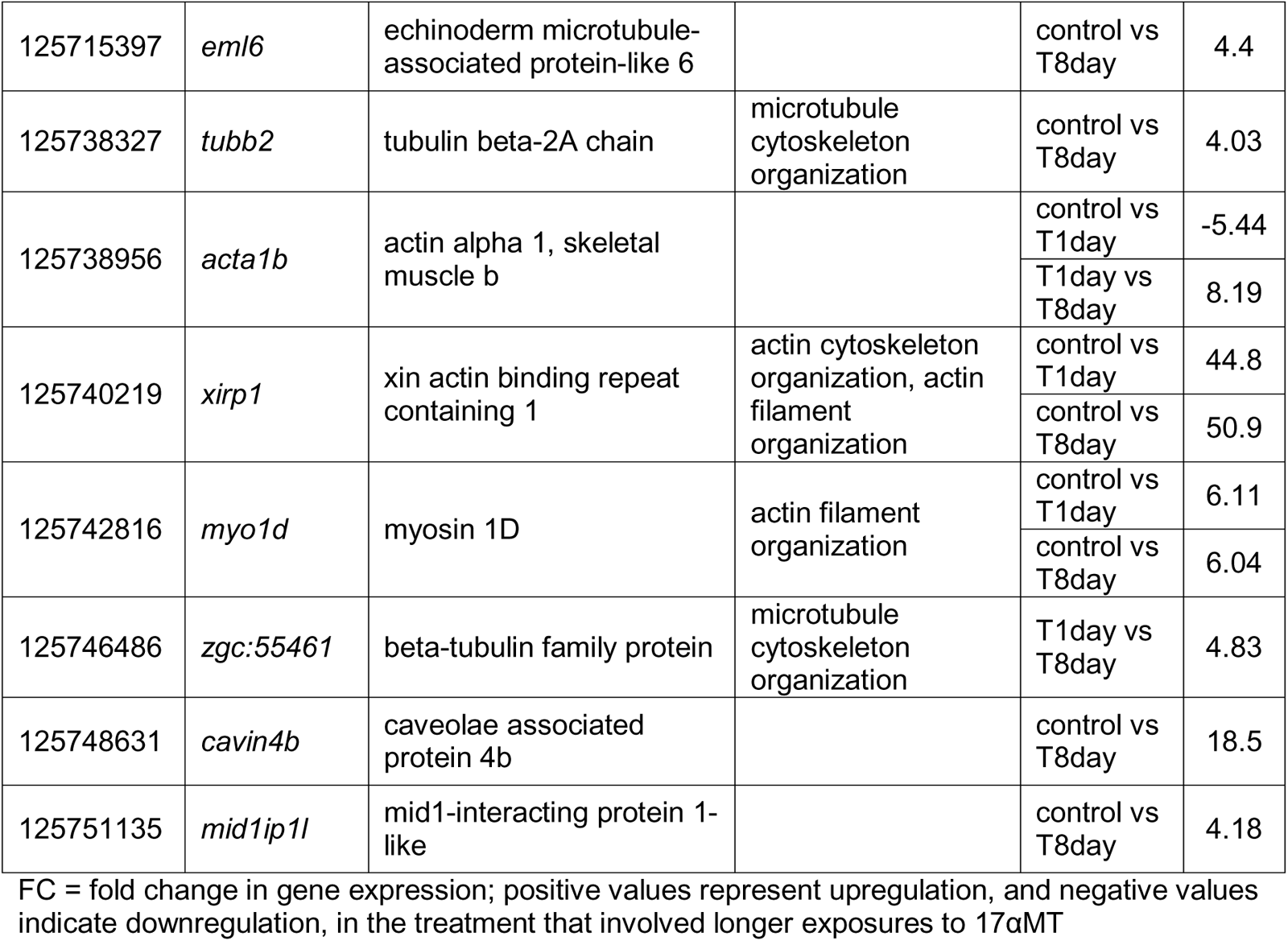
Select differentially expressed genes of highest interest likely to influence EOD duration from the general theme “cytoskeletal & sarcomeric”.

## Discussion

In this study, we sought to identify gene expression patterns related to changes in EOD duration in samples from the same species, *Brienomyrus brachyistius*. In contrast with typical studies, a single focal species reduces confounding variability introduced by interspecific differences. We experimentally treated individually housed specimens with 17αMT for 1 or 8 days and compared patterns of gene expression to individuals treated with ethanol vehicle. By leveraging on differential expression and functional enrichment analyses, we were able to systematically identify specific genes and broad functional themes that changed expression as the EOD elongated.

### EOD duration and gene expression in the electric organ

By focusing only on adult *Brienomyrus brachyistius,* our experimental design addressed confounding sources of variability such as phylogenetic divergence and age. Unlike other features of EODs like complexity or polarity, duration can change relatively quickly depending on environmental conditions. Therefore, we housed fish individually, carefully monitored their acclimation, minimized disturbances, and elicited EOD elongation with a proven hormone manipulation. The resulting EODs were consistent through time and of similar duration across treatments before addition of 17αMT, and upon its application the recipient fish steadily increased their EODs (Fig. 2, Additional file 1). This response was not evident by day 1, but it increased progressively and became striking by day 8. The critical EOD duration comparison between treatments is that of the last day of treatment (Fig. 2A, right pane) because the underlying recordings were taken shortly before tissue dissection. In other words, all the gene expression data derives from the electrocytes that generated these, and only these, EODs.

As a result of this design, we observed relatively low levels of differential expression between treatment groups (<1.17%, Table 1) and high correlation levels of gene expression across all 20 samples. Importantly, these correlation values were higher among samples of the same treatment (Additional file 2), implying that the differences in gene expression are driven by experimental manipulation. Notably, at treatment level, overall gene expression was more similar between T1day and T8day (Table 1, Fig. 3, Additional file 2), despite the seven day difference between them, compared to a one day difference between treatments control and T1day. This strongly suggests that the majority of changes in gene expression had materialized by day 1, despite little change in EOD duration at this time (Fig. 2, Additional file 1). This emphasizes the importance of including treatment T1day in our analysis: it provides granularity to the understanding of how 17αMT induces EOD elongation (Fig. 2B). In sum, given the robustness of our EOD duration and gene expression data, we expect that gene expression changes that underlie differences in EOD duration are amongst the DEG we detected.

### Functional enrichment analysis

Functional enrichment analysis greatly simplifies the interpretation of the copious amount of data generated by omic technologies like RNA sequencing, by identifying previously defined groups of related genes that differ between the experimental treatments (García-Campos et al. 2015). We employed *mitch*, a multi-contrast enrichment analysis tool, to detect gene ontology terms of biological processes jointly enriched across our three pairwise treatment contrasts (Fig. 3). To facilitate inferences about how these results may relate to our experimental manipulations and the ensuing changes in EOD duration, we further grouped these significant gene sets into broad categories (Fig. 3, Additional file 5). The per-category expression patterns uncovered a broad but coherent picture of sequential changes in gene expression that presumably start with 17αMT-induced modifications and likely culminate in a distinct set of gene expression changes that alter the EOD phenotype. We note that this analysis is limited by errors in the *D. rerio* gmt file, in the *B. brachyistius* genome annotation, and in our procedure to match *D. rerio* and *B. brachyistius* genes.

### Early Changes in Response to 17-αMT

Our data demonstrates that exposure to 17αMT triggers a signal that is transduced to downstream effectors through cell signaling pathways. Anticipated as part of the cells’ immediate response are changes in gene expression and increases in general protein-related processes, regardless of the ultimate, phenotypic effects induced. Accordingly, we broadly observed fast enrichment changes in the categories of “signaling and transduction”, “transcription and translation”, “cellular transport”, “protein catabolism”, and in the unclassified gene set “protein folding” (Fig. 3, control vs T1day contrast). After day 1, the enrichment change in these categories is generally more subdued, suggesting that they mainly represent an immediate or general reaction to hormone exposure (Fig. 3, T1day vs T8day comparison). We propose that these proximate responses ultimately induce expression changes in the genes we deem of highest interest for EOD duration, and that these genes are the effectors of the observed EOD elongation. Finally, the fast enrichment changes in the gene sets “mitochondrion organization”, “proton transmembrane transport”, and “tricarboxylic acid cycle” likely reflect increases in energy consumption, presumably a consequence of the energetic costs of the initial cellular response.

### Late Changes in Response to 17-αMT

Energy consumption, as reflected in the mentioned gene sets, consistently displays strong upregulation throughout the experiment. Suitably, longer EODs are predicted to be metabolically more costly (Hopkins 1999a), and this has been corroborated in the gymnotiform *Brachyhypopomus gauderio* (Salazar and Stoddard 2008). Furthermore, genes set in the categories “cytoskeletal and ECM organization”, “lipid metabolism”, and the unclassified gene set “muscle contraction” generally showed marked enrichment changes after day 1 (Fig. 3), and they embody several of the major features thought to be responsible for EOD variation. The genes that underlie these categories could alter the duration of the EOD by inducing changes in the surface area of the electrocyte membrane, a character that has been correlated with longer EODs in descriptive (Bennett 1971; Paul et al. 2015; Nguyen et al. 2020) and 17αMT-mediated experimental work (Bass et al. 1986; Bass and Volman 1987; Freedman et al. 1989). In addition, these genes may regulate electrocyte interactions with the extracellular matrix (ECM) and the connective tissue sheath, which may contribute to EOD phenotype (Losilla et al. 2020). Noteworthy, signal transduction through Rho GTPases (Fig. 3) is a key regulatory mechanism of the actin cytoskeleton, membrane protrusions, and adhesion complexes (Hall 1998).

Finally, the gene set “regulation of membrane potential” also exhibited a strong enrichment change after day 1. This gene set falls into our functional theme “cation homeostasis”, the remaining theme expected to influence EOD variation. Genes in this theme could alter the duration of the EOD by regulating plasma membrane ion channels, as has been demonstrated in gymnotiforms (reviewed in (Markham 2013; Dunlap et al. 2017)) and has been explored in mormyrids (Lamanna et al. 2015; Nagel et al. 2017; Swapna et al. 2018). Although our analysis only recovered one gene from this gene set (*gabrb3*, a ligand-gated chloride channel and part of the GABA receptor, Table 4), we detected voltage-gated ion channel genes differentially expressed in our contrasts and included them in our list of genes of highest interest.

### Differentially expressed genes of highest interest for EOD duration

We refined our list of candidate genes most likely to modulate EOD duration by 1) taking the genes common to the DGE analysis and to the select gene sets from the functional enrichment analysis; and, given the limitations of the functional enrichment procedure, 2) by further selecting DEG that align with previous knowledge. One bias of the latter is our inevitably incomplete understanding of the functional effects of every gene. Readers curious about the remaining DEG can find them in Additional file 3. Unless otherwise indicated, gene descriptions come from Uniprot.

### Genes that affect electrocyte morphology

The anterior and posterior faces of the electrocyte plasma membrane often increase their surface area by evaginations and invaginations of the membrane (Schwartz et al. 1975). These projections, depending on their characteristics, are referred to in the literature as *papillae*, *tubules*, *calveoli*, or *canaliculi*. The larger membrane projections are supported by the cytoskeleton (Korniienko et al. 2021). Increases in the membrane surface area are associated with longer EODs (Bennett 1971; Paul et al. 2015), mainly through projections on the anterior face (Bass et al. 1986; Freedman et al. 1989; Paul et al. 2015; Nguyen et al. 2020). These increases in membrane surface area may increase membrane capacitance that thus delays spike initiation (Bass et al. 1986; Bass and Volman 1987). Given this framework, we anticipate cytoskeletal & sarcomeric genes involved in longer EODs to be mostly upregulated in the treatment T8day, and indeed genes from this theme display this expression pattern. For example, we detected strong increases in the actin or actin-related genes: *actn1, actc1c, myh6, acta1b, xirp1,* and *myo1d*; and in the microtubule-related genes: *eml6, tubb2, zgc:55461, cavin4b,* and *mid1ip1l* (Table 2). Furthermore, we expect that this increase in surface area also requires rearrangements in the phospholipid bilayer and in the electrocyte’s interactions with the ECM and the connective tissue sheath. Previous research supports that these themes are related to variation in EOD phenotypes (lipid metabolism (Lamanna et al. 2015; Losilla et al. 2020), ECM (Losilla et al. 2020)). In fish with longer EODs, we identified increased gene expression of *elovl7a* and *selenoi,* two genes that participate in the production of membrane lipids; and of the gene *fam126a*, which plays a key role in certain cell types with expanded plasma membrane (Table 3). Likewise, the same expression pattern was observed for a collagen gene (*si:dkey-61l1.4*) and for several adhesion genes, including *thbs4b* and *cdh11*; whereas we found reduced expression of *epdl2* in these fish (Table 3). In *Paramormyrops,* we have previously identified *epdl2*, whose product may participate in cell-matrix adhesion, as a gene that could affect the EOD waveform (Losilla et al. 2020) and that underwent duplications and functional specialization (Losilla and Gallant 2023).

**Table 3.**
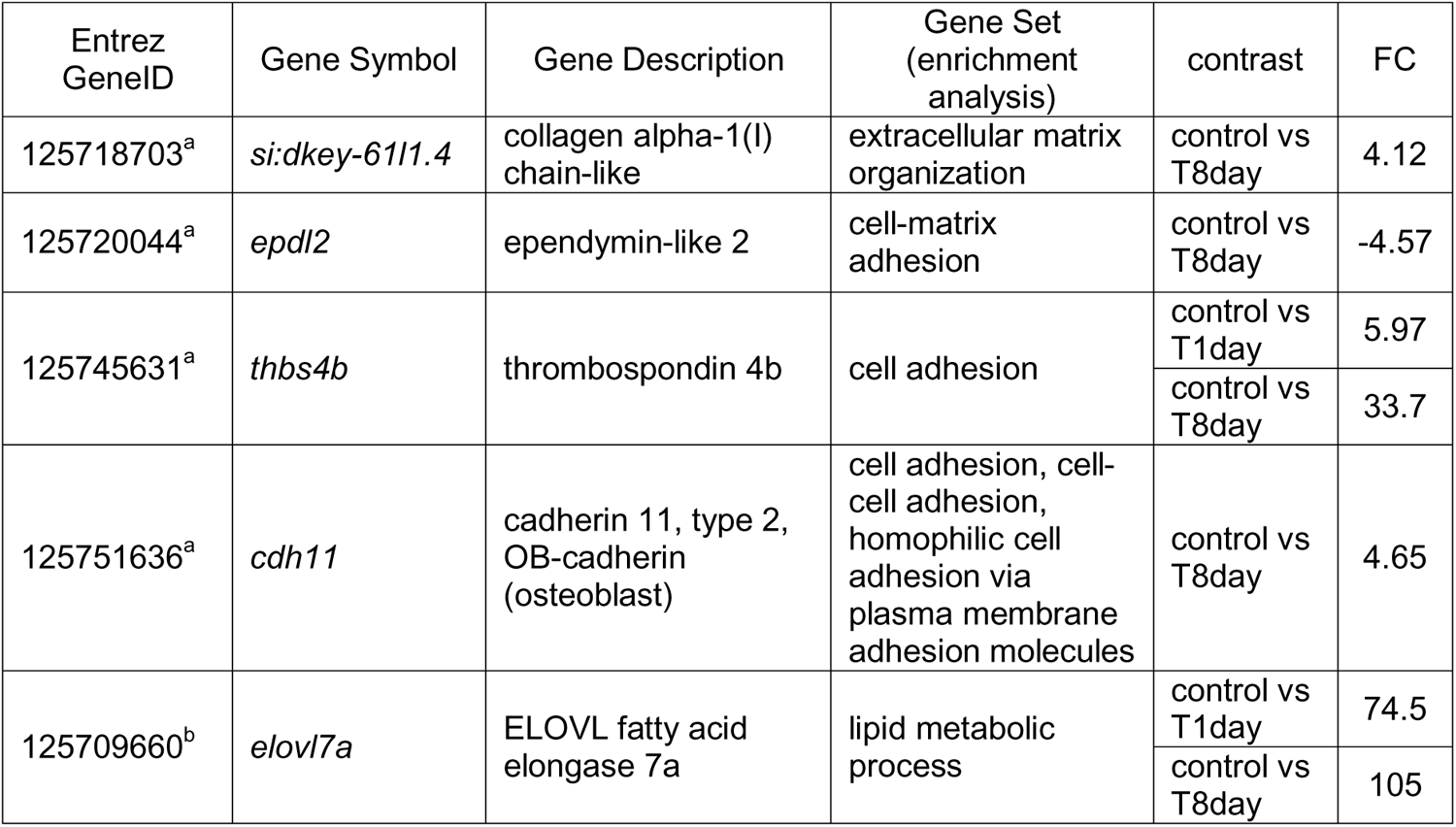
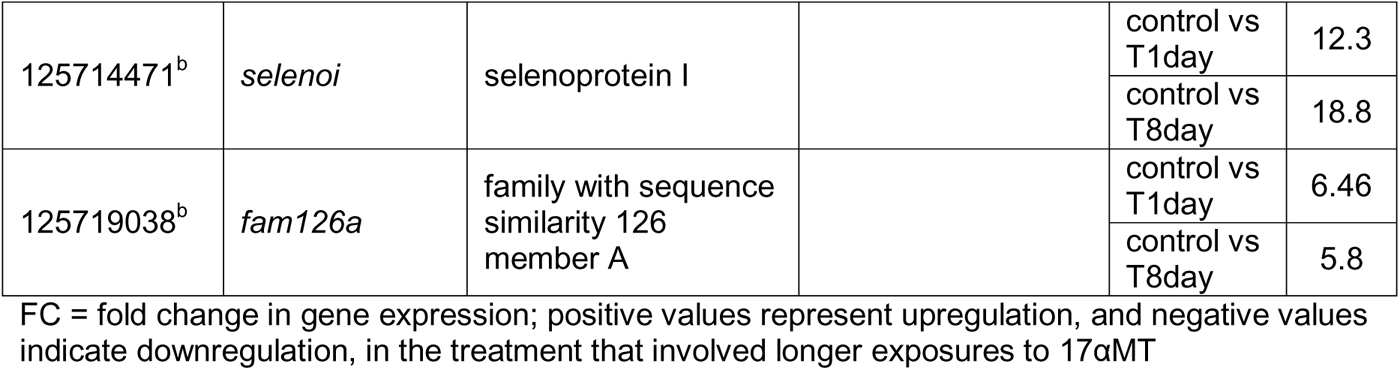
Select differentially expressed genes of highest interest likely to influence EOD duration from the general themes “extracellular matrix^a^” and “lipid metabolism^b^”.

### Genes that affect electrocyte physiology

Another mechanism to change EOD duration is to alter the electrocyte ionic currents, which can be achieved by modifying the functional properties or levels of expression of ion channels. In gymnotiforms, ionic currents are largely driven by voltage-gated Na^+^ and K^+^ currents (e.g. (Shenkel and Sigworth 1991; Ferrari and Zakon 1993)) and their modulation via androgen hormones affects EOD duration (e.g. (Zakon et al. 1991; Ferrari et al. 1995)). To the best of our knowledge, technical difficulties have hindered recordings of mormyrid electrocyte ionic currents, yet significant convergent evolution is expected. It has been proven that both Na^+^ (Zakon et al. 2006; Arnegard et al. 2010b) and K^+^ (Nagel et al. 2017; Swapna et al. 2018) voltage-gated channels are expressed in mormyrid electrocytes, and differential expression of ion channel genes has been linked with EOD variation (Lamanna et al. 2015; Losilla et al. 2020).

First, we identified a gene that codes for a Na^+^ channel beta subunit, *scn4b*, downregulated in samples with longer EODs (Table 4). This mirrors results observed in the gymnotiform *Sternopygus macrurus*, where exposure to an androgen hormone elongated the EOD and downregulated the expression of a Na^+^ channel β1-subunit, that probably speeds inactivation rates of the α subunits (Liu et al. 2007). Given this mode of action, suppression of the β subunit would likely extend the electrocyte Na^+^ currents and the EOD.

**Table 4.** Select differentially expressed genes of highest interest likely to influence EOD duration from the general theme “cation homeostasis”.

| Entrez GeneID | Gene Symbol | Gene Description | Gene Set (enrichment analysis) | contrast | FC |
| --- | --- | --- | --- | --- | --- |
| 125706737 | *kcng4a* | potassium voltage-gated channel, subfamily G, member 4a |  | control vs T8day | 7.82 |
| 125712673 | *scn4b* | sodium channel subunit beta-4-like |  | control vs T8day | -9.23 |
| 125721870 | *atp1b2b* | ATPase Na<sup>+</sup>/K<sup>+</sup> transporting subunit beta 2b |  | control vs T1day | 8.52 |
|  |  |  |  | control vs T8day | 17.9 |
| 125722963 | *gabrb3* | gamma-aminobutyric acid type A receptor subunit beta3 | regulation of membrane potential | T1day vs T8day | 7.86 |
|  |  |  |  | control vs T8day | 5.87 |
| 125738950 | *kcnc2* | potassium voltage-gated channel subfamily C member 2-like |  | control vs T1day | -4.12 |
|  |  |  |  | T1day vs T8day | -4.99 |
|  |  |  |  | control vs T8day | -20.9 |
| 125742760 | *kcnc1* | potassium voltage-gated channel subfamily C member 1-like |  | control vs T8day | -10.4 |
| 125743020 | *kcna7a1* | Potassium voltage-gated channel subfamily A member 7-like |  | control vs T8day | -2.83 |
| 125750347 | *kcna1* | potassium voltage-gated channel subfamily A member 1 |  | control vs T8day | -16.1 |

FC = fold change in gene expression; positive values represent upregulation, and negative values indicate downregulation, in the treatment that involved longer exposures to 17αMT

Second, we observed a striking coordinated regulation of at least five K^+^ voltage-gated channels genes: *kcna1, kcnc1, kcnc2, kcng4a* (Table 4); and *kcna7a1* (Entrez GeneID 125743020, potassium voltage-gated channel subfamily A member 7-like, see below). Based on the model of EOD production, APs from each electrocyte face, and therefore the resulting EOD, would elongate if K^+^ efflux is delayed or its rate reduced. Allowing K^+^ ions out of the cell is the canonical function of delayed-rectifier K^+^ channels; and two of these, Kv1.1a and Kv1.2a, are less expressed in longer EODs in *S. macrurus*, a response that can be elicited by androgen hormones (Few and Zakon 2007). We detected four differentially expressed delayed-rectifier K^+^ channels, and in agreement with expectations: all of them were strongly downregulated in fish with longer EODs (T8day treatment): *kcna1* (Kv1.1), *kcna7a1* (Kv1.7), *kcnc1* (Kv3.1), and *kcnc2* (Kv3.2). Interestingly, Kv3.2 channels show adaptations for fast repolarization in neurons (Rudy and McBain 2001; Lien and Jonas 2003), positioning *kcnc1* and *kcnc2* as candidates for electrocyte specialization in EODs of short duration. We also identified gene *kcng4a* (Kv6.4), but unlike the previous K^+^ voltage-gated channels genes, this one was found upregulated in longer EODs. The protein coded by this gene is classified as a modifier/silencer channel that modulates the delayed-rectifier K^+^ channel *kcnb1* (Kv2.1). The addition of Kv6.4 causes a shift towards more negative potentials in the voltage dependence of steady-state inactivation (Bocksteins et al. 2012; Lee et al. 2020). If this shift influences EODs, the anticipated effect would be to close the channel faster (at more negative potentials), thus reducing potassium efflux, extending the AP, and the ensuing EOD.

Finally, we note that *kcna7a* was not detected in our differential expression analysis; however, it has been reported that this gene codes for mormyrid-exclusive amino acid substitutions that enable ultra-brief EODs and that it is expressed at much higher levels than other K^+^ voltage-gated channels in mormyrid electrocytes (Swapna et al. 2018). This prompted us to take a closer look at this gene. We found four *kcna7a* genes in the *B. brachyistius* genome assembly, but we suspect that they represent two *kcna7a* paralogs and an assembly error. We call these paralogs *kcna7a1* (Entrez GeneID 125743020, and the spuriously assembled 125742764) and *kcna7a2* (Entrez GeneID 125743019, and the spuriously assembled 125742763). We conclude that *kcna7a2* is the kcna7a gene studied by (Swapna et al. 2018), because we found it expressed at much higher levels than the other K^+^ voltage-gated channels. This gene’s expression did not significantly change in our treatments. On the other hand, we found the less expressed paralog*, kcna7a1*, downregulated in T8day fish compared to control fish (FC = -2.83, FDR-corrected p-value < 0.001). Given this, we include this gene in our discussion and suggest that the significance threshold for differential expression of FC = 4 may be a too stringent for ion channel genes.

Following our findings, we propose that, in *B. brachyistius* EODs, Kv1.7a2 is the delayed-rectifier K^+^ channel that carries the most K^+^ out of the electrocytes; yet K^+^ efflux during EOD elongation is modulated, at least in part, through the concerted regulation of several, less expressed K^+^ voltage-gated channels. On the other hand, because *scn4b* is the only Na^+^ voltage-gated channel gene we detected, we propose that its regulation is the main mechanism of modulating EOD duration that involves Na^+^ flow in this species.

## Conclusion

We believe that this work represents the best controlled effort to date to study changes in gene expression that stem from changes in a single EOD character. Our results provide insights into the stepwise cellular process induced by androgen hormones and that lead to EOD elongation. A fast, coordinated response, mediated through multiple signaling pathways, activates gene expression, protein metabolism, and processes that increase energy availability. The downstream result is changes in gene expression that modify the morphological and physiological properties of electrocytes, in ways that align perfectly with theoretical expectations on how these changes could increase EOD duration. Increases in electrocyte membrane surface area are associated with longer EODs, and we detected a concordant upregulation of genes that participate in plasma membrane expansions. Importantly, our results deepen our understanding of how an increase in membrane area is accomplished, by pointing out that these morphological changes likely occur in tandem with changes to electrocyte interactions with the ECM and to the supporting cytoskeleton, including increases in specific actin and tubulin proteins, two of its essential building blocks. Similarly, our results also support that changes to the physiological properties of the electrocyte, mediated by expression changes of ion channels, influence EOD duration. They implicate changes to ionic currents that agree with current theories of EOD function and are novel examples of convergence between gymnotiforms and mormyrids. These are the downregulation in longer EODs of a Na^+^ channel beta subunit and delayed-rectifier K^+^ channels. We found multiple K^+^ voltage-gated channels, mostly but not exclusively delayed-rectifiers, consistently modulated in accordance with expectations of how they would influence EOD duration. Interestingly, these do not include *kcna7a2*, the most expressed K^+^ voltage-gated channel in electrocytes. Our results not only support and deepen current ideas of how mormyrids module EOD duration, by they also provide a list of specific genes likely to mediate this regulation. Given the central role of EOD duration to EOD diversification, and the importance of the EOD in species divergence, these genes may be significant participants in the speciation process amid African weakly electric fish.

## Notes

### Competing Interest Statement

The authors have declared no competing interest.

## References

Arnegard, M. E., S. M. Bogdanowicz, and C. D. Hopkins. 2005. Multiple cases of striking genetic similarity between alternate electric fish signal morphs in sympatry. Evolution 59:324–343.

Arnegard, M. E., and C. D. Hopkins. 2003. Electric signal variation among seven blunt-snouted Brienomyrus species (Teleostei: Mormyridae) from a riverine species flock in Gabon, Central Africa. Environ Biol Fishes 67:321–339.

Arnegard, M. E., B. S. Jackson, and C. D. Hopkins. 2006. Time-domain signal divergence and discrimination without receptor modification in sympatric morphs of electric fishes. J Exp Biol 209:2182–2198.

Arnegard, M. E., P. B. McIntyre, L. J. Harmon, M. L. Zelditch, W. G. R. Crampton, J. K. Davis, J. P. Sullivan, S. Lavoué, and C. D. Hopkins. 2010a. Sexual signal evolution outpaces ecological divergence during electric fish species radiation. Am Nat 176:335–356.

Arnegard, M. E., D. J. Zwickl, Y. Lu, and H. H. Zakon. 2010b. Old gene duplication facilitates origin and diversification of an innovative communication system--twice. Proc Natl Acad Sci U S A 107:22172– 22177.

Ashburner, M., C. A. Ball, J. A. Blake, D. Botstein, H. Butler, J. M. Cherry, A. P. Davis, K. Dolinski, S. S. Dwight, J. T. Eppig, M. A. Harris, D. P. Hill, L. Issel-Tarver, A. Kasarskis, S. Lewis, J. C. Matese, J. E. Richardson, M. Ringwald, G. M. Rubin, and G. Sherlock. 2000. Gene Ontology: tool for the unification of biology. Nat Genet 25:25–29.

Bass, A. H. 1986. Species differences in electric organs of mormyrids: Substrates for species-typical electric organ discharge waveforms. Journal of Comparative Neurology 244:313–330.

Bass, A. H., J.-P. Denizot, and M. A. Marchaterre. 1986. Ultrastructural features and hormone-dependent sex differences of mormyrid electric organs. Journal of Comparative Neurology 254:511–528.

Bass, A. H., and C. D. Hopkins. 1985. Hormonal control of sex differences in the electric organ discharge (EOD) of mormyrid fishes. Journal of Comparative Physiology A 156:587–604.

Bass, A. H., and C. D. Hopkins. 1983. Hormonal control of sexual differentiation: changes in electric organ discharge waveform. Science (1979) 220:971–974.

Bass, A. H., and S. F. Volman. 1987. From behavior to membranes: testosterone-induced changes in action potential duration in electric organs. Proceedings of the National Academy of Sciences 84:9295–9298.

Bennett, M. V. L. 1971. Electric Organs. Pp. 347–491 in W. S. Hoar and D. J. Randall, eds. Fish Physiology. Academic Press, London.

Bennett, M. V. L., and H. Grundfest. 1961. Studies on the morphology and electrophysiology of electric organs. III. Electrophysiology of electric organs in mormyrids. Pp. 113–35 in C. Chagas and A. de Carvalho, eds. Bioelectrogenesis. Elsevier, Amsterdam.

Bocksteins, E., A. J. Labro, D. J. Snyders, and D. P. Mohapatra. 2012. The Electrically Silent Kv6.4 Subunit Confers Hyperpolarized Gating Charge Movement in Kv2.1/Kv6.4 Heterotetrameric Channels. PLoS One 7:e37143.

Bolger, A. M., M. Lohse, and B. Usadel. 2014. Trimmomatic: a flexible trimmer for Illumina sequence data. Bioinformatics 30:2114–2120.

Camacho, C., G. Coulouris, V. Avagyan, N. Ma, J. Papadopoulos, K. Bealer, and T. L. Madden. 2009. BLAST+: architecture and applications. BMC Bioinformatics 10:421. BioMed Central.

Carbon, S., E. Douglass, B. M. Good, D. R. Unni, N. L. Harris, C. J. Mungall, S. Basu, R. L. Chisholm, R. J. Dodson, E. Hartline, P. Fey, P. D. Thomas, L. P. Albou, D. Ebert, M. J. Kesling, H. Mi, A. Muruganujan, X. Huang, T. Mushayahama, S. A. LaBonte, D. A. Siegele, G. Antonazzo, H. Attrill, N. H. Brown, P. Garapati, S. J. Marygold, V. Trovisco, G. dos Santos, K. Falls, C. Tabone, P. Zhou, J. L. Goodman, V. B. Strelets, J. Thurmond, P. Garmiri, R. Ishtiaq, M. Rodríguez-López, M. L. Acencio, M. Kuiper, A. Lægreid, C. Logie, R. C. Lovering, B. Kramarz, S. C. C. Saverimuttu, S. M. Pinheiro, H. Gunn, R. Su, K. E. Thurlow, M. Chibucos, M. Giglio, S. Nadendla, J. Munro, R. Jackson, M. J. Duesbury, N. Del-Toro, B. H. M. Meldal, K. Paneerselvam, L. Perfetto, P. Porras, S. Orchard, A. Shrivastava, H. Y. Chang, R. D. Finn, A. L. Mitchell, N. D. Rawlings, L. Richardson, A. Sangrador-Vegas, J. A. Blake, K. R. Christie, M. E. Dolan, H. J. Drabkin, D. P. Hill, L. Ni, D. M. Sitnikov, M. A. Harris, S. G. Oliver, K. Rutherford, V. Wood, J. Hayles, J. Bähler, E. R. Bolton, J. L. de Pons, M. R. Dwinell, G. T. Hayman, M. L. Kaldunski, A. E. Kwitek, S. J. F. Laulederkind, C. Plasterer, M. A. Tutaj, M. Vedi, S. J. Wang, P. D’Eustachio, L. Matthews, J. P. Balhoff, S. A. Aleksander, M. J. Alexander, J. M. Cherry, S. R. Engel, F. Gondwe, K. Karra, S. R. Miyasato, R. S. Nash, M. Simison, M. S. Skrzypek, S. Weng, E. D. Wong, M. Feuermann, P. Gaudet, A. Morgat, E. Bakker, T. Z. Berardini, L. Reiser, S. Subramaniam, E. Huala, C. N. Arighi, A. Auchincloss, K. Axelsen, G. Argoud-Puy, A. Bateman, M. C. Blatter, E. Boutet, E. Bowler, L. Breuza, A. Bridge, R. Britto, H. Bye-A-Jee, C. C. Casas, E. Coudert, P. Denny, A. Es-Treicher, M. L. Famiglietti, G. Georghiou, A. N. Gos, N. Gruaz-Gumowski, E. Hatton-Ellis, C. Hulo, A. Ignatchenko, F. Jungo, K. Laiho, P. le Mercier, D. Lieberherr, A. Lock, Y. Lussi, A. MacDougall, M. Ma-Grane, M. J. Martin, P. Masson, D. A. Natale, N. Hyka-Nouspikel, S. Orchard, I. Pedruzzi, L. Pourcel, S. Poux, S. Pundir, C. Rivoire, E. Speretta, S. Sundaram, N. Tyagi, K. Warner, R. Zaru, C. H. Wu, A. D. Diehl, J. N. Chan, C. Grove, R. Y. N. Lee, H. M. Muller, D. Raciti, K. van Auken, P. W. Sternberg, M. Berriman, M. Paulini, K. Howe, S. Gao, A. Wright, L. Stein, D. G. Howe, S. Toro, M. Westerfield, P. Jaiswal, L. Cooper, and J. Elser. 2021. The Gene Ontology resource: Enriching a GOld mine. Nucleic Acids Res 49:D325–D334. Oxford University Press.

Carbon, S., A. Ireland, C. J. Mungall, S. Shu, B. Marshall, S. Lewis, J. Lomax, C. Mungall, B. Hitz, R. Balakrishnan, M. Dolan, V. Wood, E. Hong, and P. Gaudet. 2009. AmiGO: Online access to ontology and annotation data. Bioinformatics 25:288–289.

Carlson, B. A. 2002. Electric signaling behavior and the mechanisms of electric organ discharge production in mormyrid fish. JOURNAL OF PHYSIOLOGY-PARIS 96:405–419. Elsevier.

Carlson, B. A., S. M. Hasan, M. Hollmann, D. B. Miller, L. J. Harmon, and M. E. Arnegard. 2011. Brain evolution triggers increased diversification of electric fishes. Science (1979) 332:583–586.

Carlson, B. A., C. D. Hopkins, and P. Thomas. 2000. Androgen correlates of socially induced changes in the electric organ discharge waveform of a mormyrid fish. Horm Behav 38:177–186.

Dunlap, K. D., A. C. Silva, G. T. Smith, and H. H. Zakon. 2017. Weakly Electric Fish: Behavior, Neurobiology, and Neuroendocrinology. Pp. 69–98 in D. W. Pfaff and M. Joëls, eds. Hormones, Brain and Behavior. Academic Press, Oxford.

Ferrari, M. B., M. L. McAnelly, and H. H. Zakon. 1995. Individual variation in and androgen-modulation of the sodium current in electric organ. Journal of Neuroscience 15:4023–4032.

Ferrari, M. B., and H. H. Zakon. 1993. Conductances contributing to the action-potential of Sternopygus electrocytes. Journal of comparative physiology A-sensory neural and behavioral physiology 173:281–292. Springer.

Few, W. P., and H. H. Zakon. 2007. Sex differences in and hormonal regulation of Kv1 potassium channel gene expression in the electric organ: Molecular control of a social signal. Dev Neurobiol 67:535–549. John Wiley & Sons.

Freedman, E. G., J. Olyarchuk, M. A. Marchaterre, and A. H. Bass. 1989. A temporal analysis of testosterone-induced changes in electric organs and electric organ discharges of mormyrid fishes. J Neurobiol 20:619–634.

Gallant, J. R., M. E. Arnegard, J. P. Sullivan, B. A. Carlson, and C. D. Hopkins. 2011. Signal variation and its morphological correlates in Paramormyrops kingsleyae provide insight into the evolution of electrogenic signal diversity in mormyrid electric fish. Journal of Comparative Physiology A 197:799–817.

Gallant, Jason. R., C. D. Hopkins, and D. L. Deitcher. 2012. Differential expression of genes and proteins between electric organ and skeletal muscle in the mormyrid electric fish Brienomyrus brachyistius. Journal of Experimental Biology 215:2479–2494.

Gallant, Jason. R., L. L. Traeger, J. D. Volkening, H. Moffett, P.-H. Chen, C. D. Novina, G. N. Phillips, R. Anand, G. B. Wells, M. Pinch, R. Guth, G. A. Unguez, J. S. Albert, H. H. Zakon, M. P. Samanta, and M. R. Sussman. 2014. Genomic basis for the convergent evolution of electric organs. Science (1979) 344:1522–1525.

García-Campos, M. A., J. Espinal-Enríquez, and E. Hernández-Lemus. 2015. Pathway analysis: State of the art. Front Physiol 6:1–16.

Grabherr, M. G., B. J. Haas, M. Yassour, J. Z. Levin, D. A. Thompson, I. Amit, X. Adiconis, L. Fan, R. Raychowdhury, Q. Zeng, Z. Chen, E. Mauceli, N. Hacohen, A. Gnirke, N. Rhind, F. di Palma, B. W. Birren, C. Nusbaum, K. Lindblad-Toh, N. Friedman, and A. Regev. 2011. Full-length transcriptome assembly from RNA-Seq data without a reference genome. Nat Biotechnol 29:644–52. Nature Publishing Group, a division of Macmillan Publishers Limited. All Rights Reserved.

Haas, B. J., A. Papanicolaou, M. Yassour, M. Grabherr, P. D. Blood, J. Bowden, M. B. Couger, D. Eccles, B. Li, M. Lieber, M. D. Macmanes, M. Ott, J. Orvis, N. Pochet, F. Strozzi, N. Weeks, R. Westerman, T. William, C. N. Dewey, R. Henschel, R. D. Leduc, N. Friedman, and A. Regev. 2013. De novo transcript sequence reconstruction from RNA-seq using the Trinity platform for reference generation and analysis. Nat Protoc 8:1494–512. Nature Publishing Group, a division of Macmillan Publishers Limited. All Rights Reserved.

Hall, A. 1998. Rho GTpases and the actin cytoskeleton. Science (1979) 279:509–514.

Herfeld, S., and P. Moller. 1998. Effects of 17α-methyltestosterone on sexually dimorphic characters in the weakly discharging electric fish, Brienomyrus niger (Gunther, 1866) (Mormyridae): Electric organ discharge, ventral body wall indentation, and anal-fin ray bone expansion. Horm Behav 34:303–319.

Hopkins, C. D. 1986. Behavior of Mormyridae. Pp. 527–576 in T. H. Bullock and W. Heiligenberg, eds. Electroreception. Wiley, New York.

Hopkins, C. D. 1999a. Design features for electric communication. Journal of Experimental Biology 202:1217–1228. Co Biol.

Hopkins, C. D. 1981. On the diversity of electric signals in a community of mormyrid electric fish in West Africa. Am Zool 21:211–222. Oxford University Press.

Hopkins, C. D. 1999b. Signal evolution in electric communication. Pp. 461– 491 in M. Hauser and M. Konishi, eds. The design of animal communication. MIT Press, Cambridge, MA.

Hopkins, C. D., and A. H. Bass. 1981. Temporal coding of species recognition signals in an electric fish. Science (1979) 212:85–87.

Kaspi, A., and M. Ziemann. 2020. Mitch: Multi-contrast pathway enrichment for multi-omics and single-cell profiling data. BMC Genomics 21:1–17. BMC Genomics.

Korniienko, Y., R. Tiedemann, M. Vater, and F. Kirschbaum. 2021. Ontogeny of the electric organ discharge and of the papillae of the electrocytes in the weakly electric fish Campylomormyrus rhynchophorus (Teleostei: Mormyridae). Journal of Comparative Neurology 529:1052–1065.

Kramer, B. 1974. Electric organ discharge interaction during interspecific agonistic behaviour in freely swimming mormyrid fish. J Comp Physiol 93:203–235. Springer-Verlag.

Lamanna, F., F. Kirschbaum, I. Waurick, C. Dieterich, and R. Tiedemann. 2015. Cross-tissue and cross-species analysis of gene expression in skeletal muscle and electric organ of African weakly-electric fish (Teleostei; Mormyridae). BMC Genomics 16:668. BMC Genomics.

Langmead, B., and S. L. Salzberg. 2012. Fast gapped-read alignment with Bowtie 2. Nat Methods 9:357–359. Nature Publishing Group.

Lavoué, S., M. E. Arnegard, J. P. Sullivan, and C. D. Hopkins. 2008. Petrocephalus of Odzala offer insights into evolutionary patterns of signal diversification in the Mormyridae, a family of weakly electrogenic fishes from Africa. J Physiol Paris 102:322–339. Elsevier.

Lee, M. C., M. S. Nahorski, J. R. F. Hockley, V. B. Lu, G. Ison, L. A. Pattison, G. Callejo, K. Stouffer, E. Fletcher, C. Brown, I. Drissi, D. Wheeler, P. Ernfors, D. Menon, F. Reimann, E. S. J. Smith, and C. G. Woods. 2020. Human Labor Pain Is Influenced by the Voltage-Gated Potassium Channel KV6.4 Subunit. Cell Rep 32. Elsevier B.V.

Li, B., and C. N. Dewey. 2011. RSEM: accurate transcript quantification from RNA-Seq data with or without a reference genome. BMC Bioinformatics 12:323. BioMed Central.

Lien, C. C., and P. Jonas. 2003. Kv3 potassium conductance is necessary and kinetically optimized for high-frequency action potential generation in hippocampal interneurons. Journal of Neuroscience 23:2058–2068.

Lissmann, H. W., and K. E. Machin. 1958. The mechanism of object location in Gymnarchus niloticus and similar fish. Journal of Experimental Biology 35:451–486.

Liu, H., M. M. Wu, and H. H. Zakon. 2007. Individual variation and hormonal modulation of a sodium channel β subunit in the electric organ correlate with variation in a social signal. Dev Neurobiol 67:1289–1304. John Wiley & Sons.

Losilla, M., and J. R. Gallant. 2023. Molecular evolution of the ependymin-related gene epdl2 in African weakly electric fish. G3 Genes|Genomes|Genetics 13. Oxford University Press (OUP).

Losilla, M., D. M. Luecke, and J. R. Gallant. 2020. The transcriptional correlates of divergent electric organ discharges in Paramormyrops electric fish. BMC Evol Biol 20:6.

Markham, M. R. 2013. Electrocyte physiology: 50 years later. Journal of Experimental Biology 216:2451– 2458.

Markham, M. R., and H. H. Zakon. 2014. Ionic Mechanisms of Microsecond-Scale Spike Timing in Single Cells. Journal of Neuroscience 34:6668–6678.

Mills, A., and H. H. Zakon. 1987. Coordination of EOD frequency and pulse duration in a weakly electric wave fish: the influence of androgens. Journal of Comparative Physiology A 161:417–430. Springer-Verlag.

Möhres, F. P. 1957. Elektrische Entladungen im Dienste der Revierabgrenzung bei Fischen. Naturwissenschaften 44:431–432. Springer-Verlag.

Nagel, R., F. Kirschbaum, and R. Tiedemann. 2017. Electric organ discharge diversification in mormyrid weakly electric fish is associated with differential expression of voltage-gated ion channel genes. J Comp Physiol A Neuroethol Sens Neural Behav Physiol 203:183–195. Springer Berlin Heidelberg.

Nguyen, L., V. Mamonekene, M. Vater, P. Bartsch, R. Tiedemann, and F. Kirschbaum. 2020. Ontogeny of electric organ and electric organ discharge in Campylomormyrus rhynchophorus (Teleostei: Mormyridae). J Comp Physiol A Neuroethol Sens Neural Behav Physiol 206:453–466. Springer Berlin Heidelberg.

Paul, C., V. Mamonekene, M. Vater, P. G. D. Feulner, J. Engelmann, R. Tiedemann, and F. Kirschbaum. 2015. Comparative histology of the adult electric organ among four species of the genus Campylomormyrus (Teleostei: Mormyridae). J Comp Physiol A Neuroethol Sens Neural Behav Physiol 201:357–374.

Rabosky, D. L., F. Santini, J. Eastman, S. A. Smith, B. Sidlauskas, J. Chang, and M. E. Alfaro. 2013. Rates of speciation and morphological evolution are correlated across the largest vertebrate radiation. Nat Commun 4:1–8.

Robinson, M. D., and A. Oshlack. 2010. A scaling normalization method for differential expression analysis of RNA-seq data. Genome Biol 11:R25.

Rudy, B., and C. J. McBain. 2001. Kv3 channels: Voltage-gated K+ channels designed for high-frequency repetitive firing. Trends Neurosci 24:517–526.

Salazar, V. L., and P. K. Stoddard. 2008. Sex differences in energetic costs explain sexual dimorphism in the circadian rhythm modulation of the electrocommunication signal of the gymnotiform fish Brachyhypopomus pinnicaudatus. J Exp Biol 211:1012–1020.

Schwartz, I. R., G. D. Pappas, and M. V. L. Bennett. 1975. The fine structure of electrocytes in weakly electric teleosts. J Neurocytol 4:87–114.

Shenkel, S., and F. J. Sigworth. 1991. Patch recordings from the electrocytes electrophorus electricus: Na currents and PNa/PK variability. Journal of General Physiology 97:1013–1041. Rockefeller Univ Press.

Stoddard, P. K., and M. R. Markham. 2008. Signal Cloaking by Electric Fish. Bioscience 58:415.

Subhash, S., and C. Kanduri. 2016. GeneSCF: A real-time based functional enrichment tool with support for multiple organisms. BMC Bioinformatics 17. BioMed Central Ltd.

Sullivan, J. P., S. Lavoué, and C. D. Hopkins. 2002. Discovery and phylogenetic analysis of a riverine species flock of African electric fishes (Mormyridae: Teleostei). Evolution (N Y) 56:597–616. Society for the Study of Evolution.

Swapna, I., A. Ghezzi, J. M. York, M. R. Markham, D. B. Halling, Y. Lu, J. R. Gallant, and H. H. Zakon. 2018. Electrostatic Tuning of a Potassium Channel in Electric Fish. Current Biology 28:2094-2102.e5. Cell Press.

Szabo, T. 1961. Les Organes Electriques des Mormyrides. Pp. 20–24 in C. Chagas and A. de Carvalho, eds. Bioelectrogenesis. Elsevier, New York.

The UniProt Consortium. 2019. UniProt: a worldwide hub of protein knowledge. Nucleic Acids Res 47:D506–D515.

von der Emde, G., M. Amey, J. Engelmann, S. Fetz, C. Folde, M. Hollmann, M. Metzen, and R. Pusch. 2008. Active electrolocation in Gnathonemus petersii: Behaviour, sensory performance, and receptor systems. J Physiol Paris 102:279–290. Elsevier Ltd.

Wu, C., I. MacLeod, and A. I. Su. 2013. BioGPS and MyGene.info: Organizing online, gene-centric information. Nucleic Acids Res 41:D561–D565.

Xin, J., A. Mark, C. Afrasiabi, G. Tsueng, M. Juchler, N. Gopal, G. S. Stupp, T. E. Putman, B. J. Ainscough, O. L. Griffith, A. Torkamani, P. L. Whetzel, C. J. Mungall, S. D. Mooney, A. I. Su, and C. Wu. 2016. High-performance web services for querying gene and variant annotation. Genome Biol 17:91. BioMed Central.

Zakon, H. H., Y. Lu, D. J. Zwickl, and D. M. Hillis. 2006. Sodium channel genes and the evolution of diversity in communication signals of electric fishes: Convergent molecular evolution. Proceedings of the National Academy of Sciences 103:3675–3680. Section of Neurobiology, University of Texas, Austin, TX 78712, USA.

Zakon, H. H., A. C. Mills, and M. B. Ferrari. 1991. Androgen-dependent modulation of the electrosensory and electromotor systems of a weakly electric fish. Seminars in Neuroscience 3:449–457.

