## Additional File 1 for "Mechanistic insights into gene expression changes and electric organ discharge elongation in mormyrid electric fish"

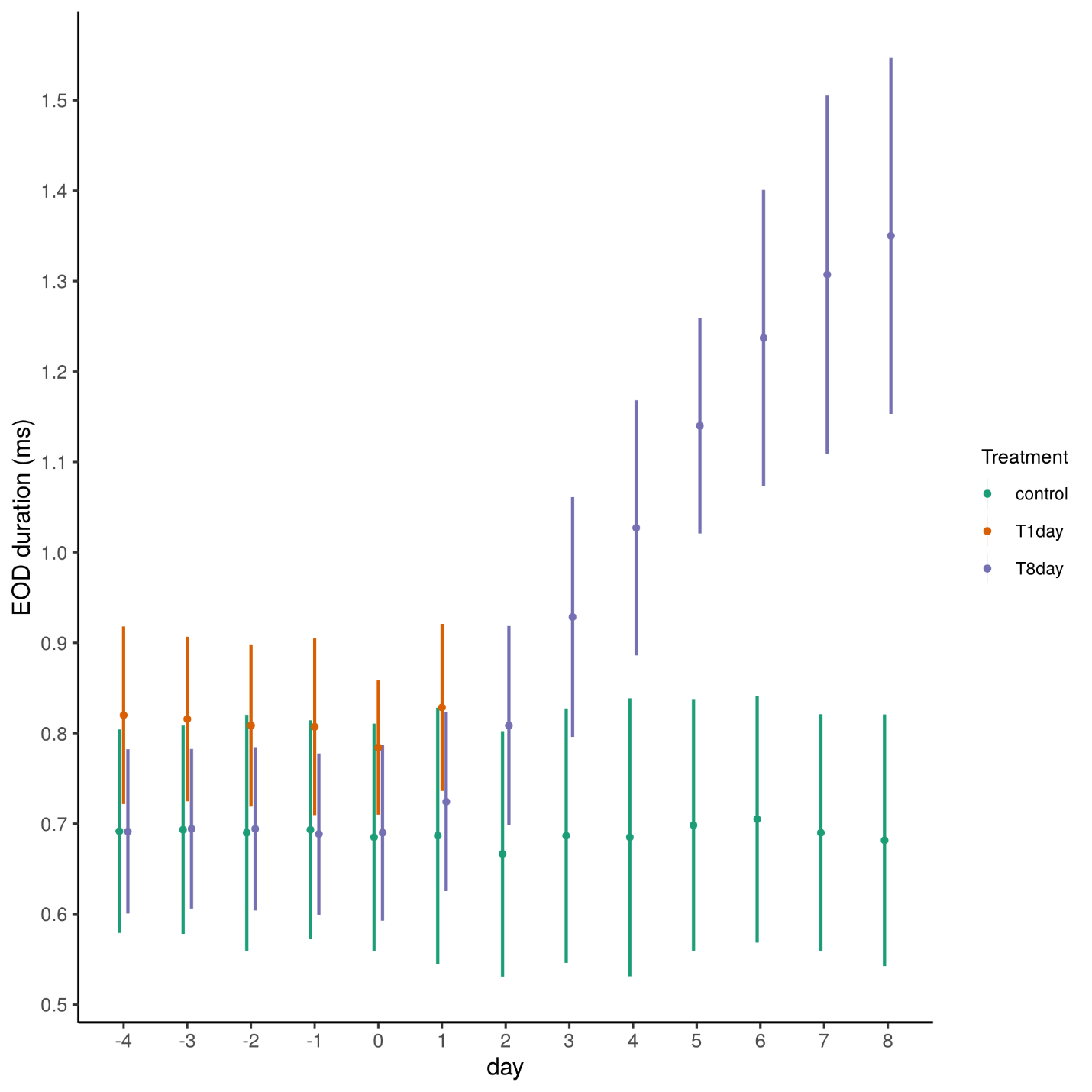

EOD duration per treatment throughout the experiment. Each colored circle and its vertical lines represent the mean EOD duration +/- standard deviation per treatment and day. A small horizontal jitter was added to better visualize overlapping values. Days -4 to 0 are part of the acclimation period, treatment-specific manipulations were performed on Day 0 after taking EOD recordings.
