## Additional File 2 for "Mechanistic insights into gene expression changes and electric organ discharge elongation in mormyrid electric fish"

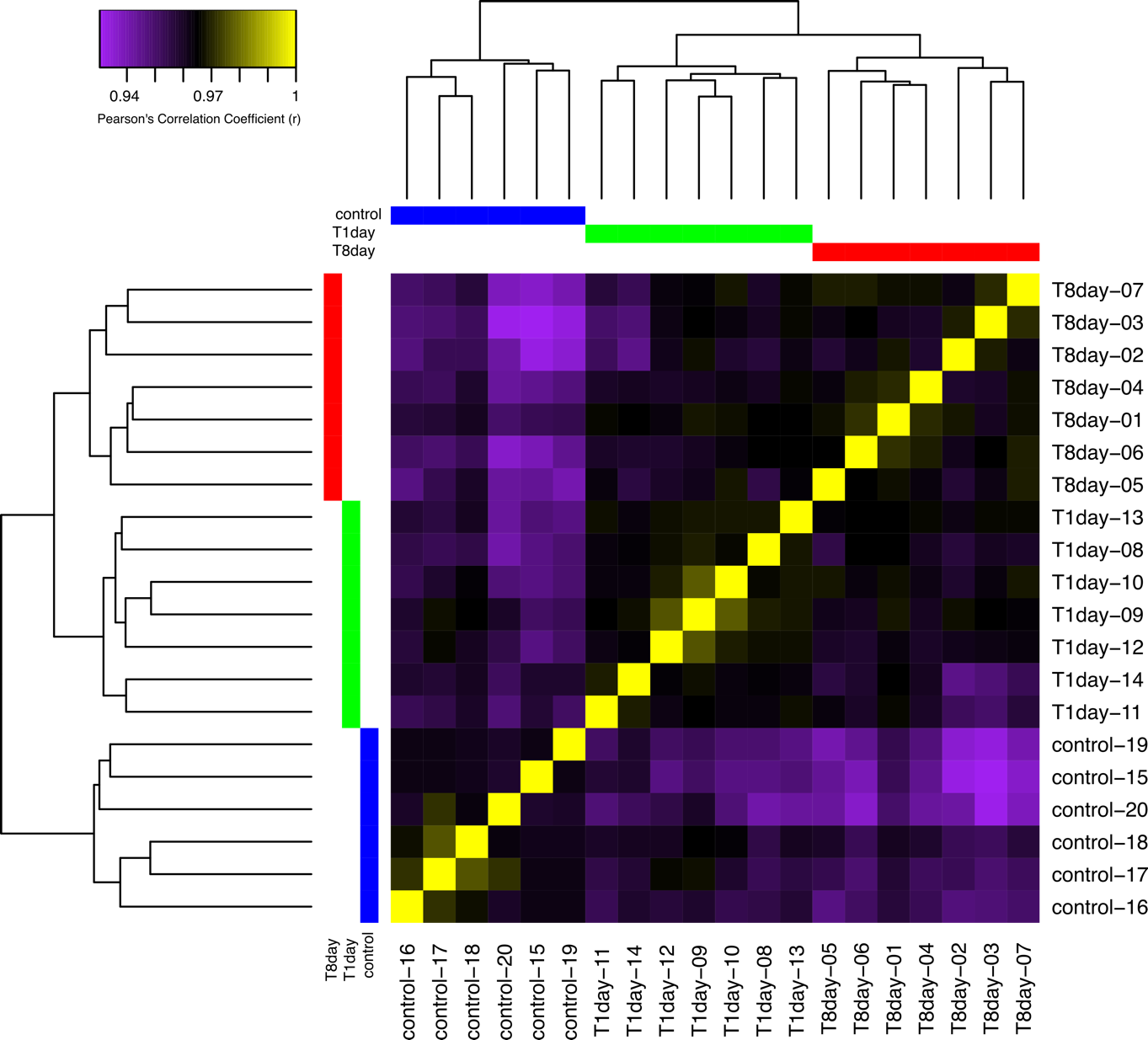

Heatmap of sample by sample correlations in gene expression, and the inferred relationships among treatments from these expression correlation values.
