## Additional File 5 for "Mechanistic insights into gene expression changes and electric organ discharge elongation in mormyrid electric fish"

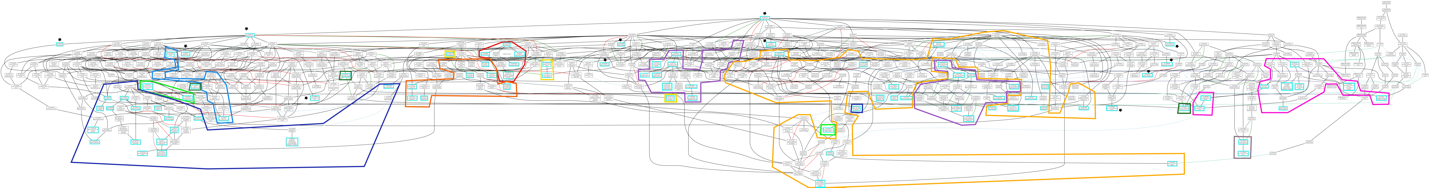

Gene Ontology graph with the 96 enriched GO terms (cyan boxes) and their grouping into 11 broad categories (outlined by the colored shapes). 12 GO terms (black stars) remained unclassified because they were too general or too isolated.
